# Benchmarking of variant pathogenicity prediction methods using a population genetics approach

**DOI:** 10.1101/2025.03.16.643565

**Authors:** Mikhail Gudkov, Loïc Thibaut, Steven Monger, Debjani Das, Congenital Heart Disease Synergy Study group, David S. Winlaw, Sally L. Dunwoodie, Eleni Giannoulatou

## Abstract

**Motivation:** Variant pathogenicity predictors are essential for identifying new associations between genetic variants and rare diseases. However, despite the availability of numerous predictors, there is no clear consensus on which methods provide the most reliable results. The common practice of training, testing, and benchmarking these predictors using known variant sets from disease or mutagenesis studies raises concerns about ascertainment bias and data circularity.

**Results:** We benchmarked commonly used pathogenicity predictors using an orthogonal approach that does not rely on predefined “ground truth” datasets. By leveraging population-level genomic data from gnomAD and the Context-Adjusted Proportion of Singletons (CAPS) metric, we identified CADD and REVEL as the best-performing predictors for distinguishing extremely deleterious variants from moderately deleterious ones. REVEL demonstrated superior calibration. Additionally, we show that CAPS can serve as a meta-analysis tool for interpreting variant annotations and highlight biases in ClinVar-based predictor training.

**Availability and Implementation:** CAPS analysis and benchmarking results are available at https://github.com/mgudVCCRI/PopGenVariantFiltering

## Introduction

The search for causal variants in the context of rare diseases is invariably complicated and disease genomics analysts often deal with a myriad of candidate variants. To filter out all candidate variants that are unlikely to contribute to the disease in question, genetic variant curators should have access to as much reliable information describing each variant as possible.

However, the problem of analysis and prioritisation of genetic single-nucleotide variants (SNVs) is hindered by the oversaturation of the disease genomics field with various pathogenicity prediction methods^1^. Even though these tools are integral to the multi-step process of variant prioritisation at large, nowadays variant analysts have to manually select among hundreds of annotations that could be available for each SNV. The main problem with this somewhat biased approach is that it may affect decision-making, for example with regards to which variants should be prioritised for further, in-depth analysis^2^.

Despite the obvious usefulness of pathogenicity predictions, most methods that produce these kinds of annotations tend to, in one way or another, employ complex, convoluted models. As a result, such methods work as “black boxes” that make predictions^2^. Furthermore, not only do these methods have different architectures, but they also use assorted sets of variant features, with some methods relying solely on DNA sequence conservation or clinical annotations, whereas others include tens of seemingly disparate variant characteristics^3^.

Several studies have been published over the last decade focusing on either the problem of variant prioritisation itself or which of the currently available predictors perform best in addressing this problem. Some studies have argued that variant pathogenicity prediction is best done at the disease^4,5^, gene^6,7^ or even protein domain^8^ levels. However, the sets of pathogenicity predictors used in currently available benchmarking studies are strikingly different, with some studies including fewer than four methods or ignoring some of the best-performing ones, such as REVEL^6,7,9–13^. More importantly, the standard way of training, testing, and benchmarking of pathogenicity predictors depends on known sets of variants from disease and mutagenesis studies, which raises issues related to ascertainment bias and data circularity^2,14–16^. Therefore, we believe that variant prioritisation remains a key challenge in rare disease genomics, despite the abundance of different pathogenicity prediction methods available, and hence, an orthogonal approach is required.

The last few years have seen an emergence of alternative and simpler methods for studying variant deleteriousness that are designed based on “snapshots” of genetic data from large databases of population-level genomic information, such as gnomAD^17,18^. Because such methods are based on well-established principles, they can provide more transparent and understandable results than some of the more complex approaches.

Negative selection can be used as a simple measure of deleterious effects of any particular group of variants. One way to estimate negative selection is to compare the number of rare variants observed in a given dataset with the expected level of rare variation. For simplicity, rare variants are often restricted to singletons (i.e., variants that are observed only once in the dataset and therefore their allele count is equal to 1). Examples of population genomics methods that utilise this approach to estimate negative selection and deleteriousness include the Mutability-Adjusted Proportion of Singletons (MAPS) metric^17^ and our recently developed Context-Adjusted Proportion of Singletons (CAPS) metric^19^, a more robust and accurate version of MAPS. MAPS and CAPS are both calculated as the difference between observed number of singletons and expected number of singletons for a group of SNVs divided by the total number of variants in that group. In MAPS, the expected number is obtained from a singletons-by-mutability model that groups all SNVs by their trinucleotide context sequence (with the affected nucleotide in the middle and one adjacent base on the left and on the right of it) to estimate mutation rates and returns an expected number of singletons, which is then summed over all contexts^20–22^. In contrast, CAPS uses per-context proportions of singletons directly to estimate the expected level, which results in a superior performance compared to MAPS^19^.

The possibility of using the Adjusted Proportion of Singletons family of metrics, and specifically MAPS, for benchmarking was first expressed in 2018, before the official publication of the currently latest version of MAPS in 2020^23^. However, to our knowledge, no studies have used such metrics to test the accuracy of pathogenicity predictors to date.

In this study, we assessed the average deleteriousness of variant sets resulting from the application of different quality control (QC) criteria using the CAPS metric. We demonstrate a proof-of-concept analysis showing how population genomics methods can aid in elucidating the performance of various individual tool-threshold pairs as well as complex combinations of QC filters. Given that the ClinVar database remains one of the most popular resources for testing, calibration, and benchmarking of variant pathogenicity predictors^3,6,7,9,10,24^, we sought to assess how well the estimates of CAPS agree with the deleteriousness of different classes of variants in this database.

## Materials and methods

We used variants from gnomAD v2.1.1 whole-exome sequencing (WES) data. The variants were preprocessed using the same steps as described in the CAPS paper^19^, which in turn are based on the preprocessing from the 2020 gnomAD flagship paper^17^. Specifically, we only retained those variants that did not have any filter flags and were called in at least 80% of potential carriers. For example, for chromosome Y variants, we excluded all variants where the call was made in less than 80% of the total number of potential male carriers, whereas for variants in autosomes, the number was 80% of two times the total number of male and female samples. We also subsequently removed all variants with coverage below 30. The resulting variant numbers are shown by class in Table S1.

We annotated missense variants from this set with deleteriousness scores from dbNSFP v4.2a^25^, taking the most deleterious prediction for each variant. Specifically, from dbNSFP we extracted the scores for REVEL v1.3, CADD v1.6, PolyPhen-2 v2.2.2, MutPred v1.2, SIFT Ensembl 66, MutationTaster v2, MutationAssessor release 3, FATHMM v2.3, MetaLR v1.0, MetaSVM v1.0 and PROVEAN v1.1. To select some commonly used tool-threshold pairs, we used a sample variant report (Figure S1) for a missense variant from the Clinical Genome Resource (ClinGen), a database of the clinical relevance of variants for precision medicine. Our ClinVar analyses, however, were not limited to missense variants. We used ClinVar version “2019-07”, available as part of gnomAD v2.1.1.

CAPS values were calculated by grouping variants of interest by their trinucleotide context (96 contexts with additional eight methylation-adjusted contexts), extracting the proportion of singletons for each context from the reference set, and taking the difference between the total number of observed singletons and the total number of expected singletons divided by the total number of all variants.

As part of the analysis of average deleteriousness of different ClinVar categories, we compared variants of uncertain significance (VUSs) assessed by the ClinVar expert panel with our in-house selection of VUSs specific to congenital heart disease (CHD) compiled by variant curators at the Victor Chang Cardiac Research Institute (VCCRI)^26^. To this end, we selected 75 prioritised VUSs from ongoing VCCRI CHD studies and retrieved these variants from gnomAD. Out of 75 variants, only 31 were present in gnomAD v2.1.1, all of which were missense variants. Since CAPS uses information about each variant’s allele count in gnomAD, we had to discard the remaining 44 variants. Importantly, only 16 of the 31 gnomAD variants in our set have any matching ClinVar records to date.

To assess the effect of applying multiple QC criteria on resulting average deleteriousness, we calculated CAPS scores for assorted variant sets using incremental addition of different filters. For example, applying the condition that all variants in a set have to pass a certain threshold based on a tool’s deleteriousness score on top of another such filter often results in a smaller variant set with a different average CAPS value. We refer to this procedure as “chaining” of filters (see Supplemental Note 1).

## Results

### CAPS as a benchmarking tool

We used our recently developed CAPS metric as a benchmarking tool, where higher values indicate stronger deleteriousness. We selected some of the most popular pathogenicity predictors and assessed how well they differentiate between moderately deleterious and extremely deleterious missense variants. Figure 1 illustrates the performance of each of the predictors in question in terms of this variant separation ability (i.e. the ability of a predictor to separate the most deleterious variants through its scores). Given that MutPred does not seem to provide a score for variants that it deems to be nondeleterious, we had to only include those variants, for which all scores were available. As shown in the figure, CADD and REVEL have the best variant separation performance in this dataset, with REVEL excelling at differentiating between variants even beyond the top ~3% point, due to its calibration. This indicates that in order to select variants with highest deleteriousness, one only needs to select the desired cut-off point (e.g. top 10%) based on the score with the best variant separation ability.

**Figure 1.**
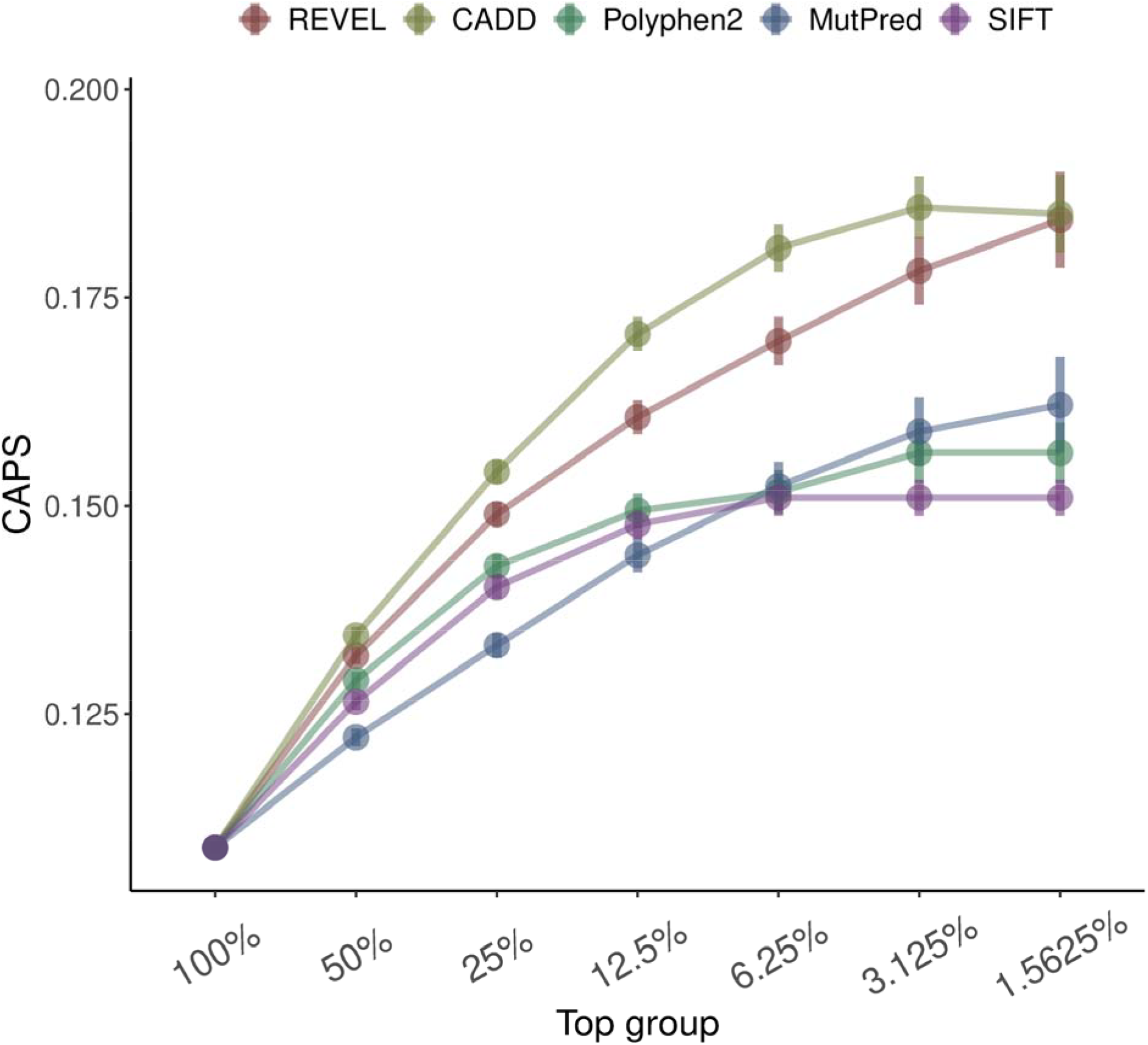
Comparison of pathogenicity predictors by their ability to separate variants with extremely high average deleteriousness from those with moderately high average deleteriousness. CAPS scores for exponential top groups of several main deleteriousness scores are shown. Error bars are 95% binomial confidence intervals. “100%”: 3259875 missense variants with all scores available (i.e. NAs removed).

### Analysis of variant filtering setups

Figure 2 shows an example of how CAPS can be used as a deleteriousness metric for threshold-based variant filtering setups. We used two sets of thresholds for three different predictors, namely SIFT, PolyPhen and CADD, with all thresholds in the first set being less stringent than in the second set. Specifically, we compared some previously suggested thresholds (SIFT <0.49, PolyPhen >0.022, CADD >10.37)^27^ with their default/recommended counterparts (SIFT <0.05, PolyPhen >0.8, CADD >20)^24,27–29^. We calculated CAPS scores for each of the resulting six groups of variants and created two additional variant sets, with only those variants that passed all three thresholds. Unsurprisingly, all eight CAPS values were higher on the negative-selection scale than the average genome-wide missense level. As expected, stricter thresholds produced a smaller subset of variants with higher levels of deleteriousness. In addition, the figure shows that when pathogenicity predictors are used in combination and multiple filtering criteria are applied, the produced variant set tends to contain variants with higher average deleteriousness than when only one method is selected.

**Figure 2.**
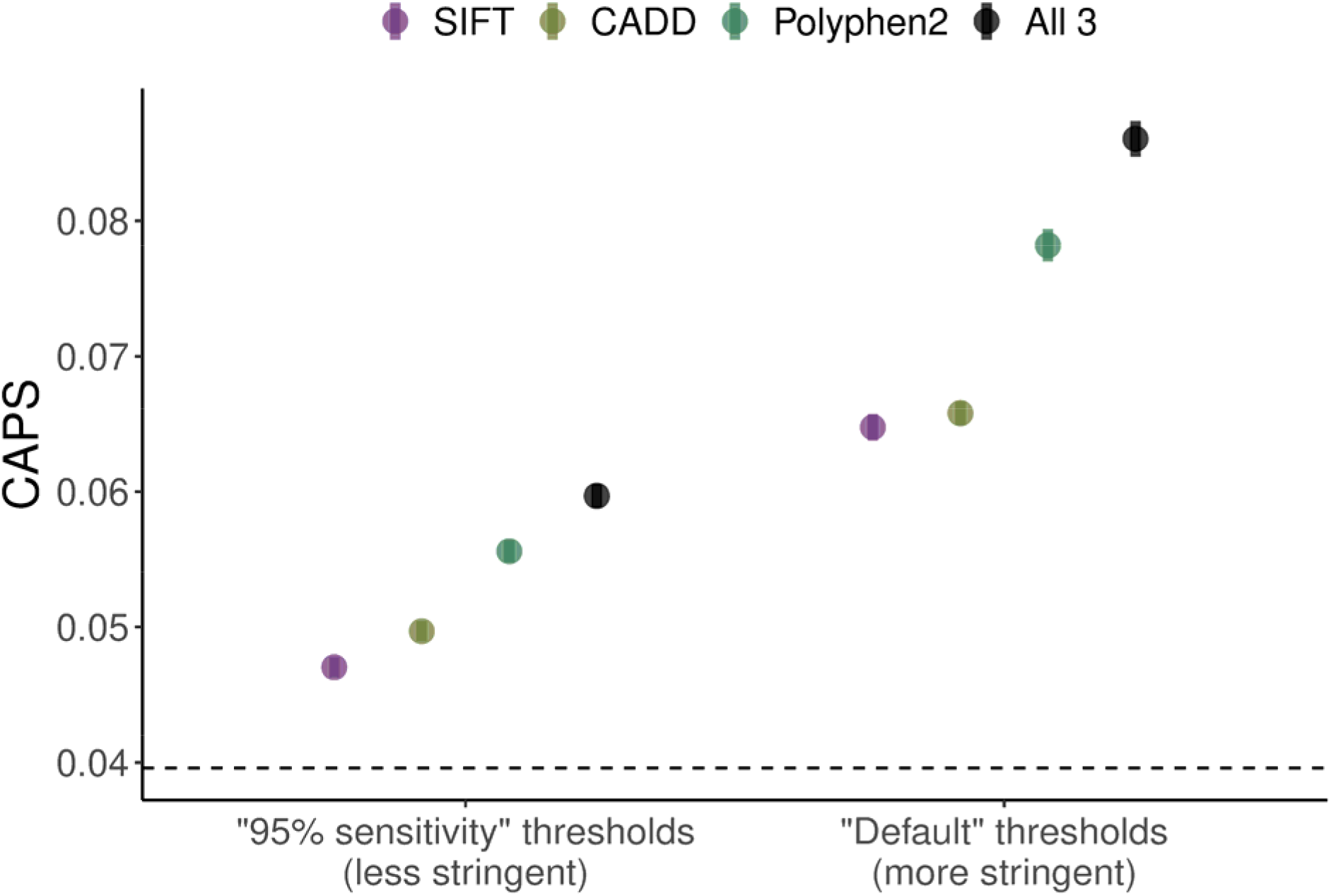
The effect of using stringent thresholds and “chaining” of filters on resulting CAPS scores. CAPS scores are shown for groups of variants that meet previously suggested thresholds for SIFT, PolyPhen and CADD. “95% sensitivity”: the less stringent thresholds from Jagadeesh et al. 2016 that were derived empirically to achieve 95% sensitivity for each tool on a test dataset; “Default”: the more stringent thresholds commonly used/recommended in literature. “All”: variants that meet all 3 thresholds. Higher values of CAPS suggest higher deleteriousness of those variants passing the more stringent filters. Dashed line indicates missense level averaged over all WES variants. Error bars are 95% binomial confidence intervals. Missense variants only. Variant numbers for each group are shown in Table S4.

### Finding critical piece of information in a variant report

CAPS can also be used to find key pieces of information in a variant report containing annotations from pathogenicity predictors. We focused on the ten annotations that were in active use in ClinGen variant reports at the time of writing and were readily available through dbNSFP for the gnomAD dataset. This selection allowed us to construct a realistic, proof-of-concept demonstration of CAPS using variant sets similar to those encountered in real-world variant interpretation workflows. The heatmap in Figure 3A shows CAPS scores for groups of variants that passed any two ClinGen thresholds. For example, selecting SIFT on the X-axis and CADD on the Y-axis corresponds with a variant set resulting from applying the ClinGen recommended threshold for SIFT in combination with the ClinGen threshold for CADD, regardless of the order in which these two QC criteria are applied. The figure shows that certain combinations of filters produce variant sets with extremely high deleteriousness. In this case report, the annotation that contributed the most to the high deleteriousness scores was the REVEL filter of 0.75 (Figure S1). Therefore, the fact that the missense variant in question passed this threshold for REVEL—and all gnomAD variants that pass this threshold are very deleterious—is the most important piece of information in this report. All other filters provide zero additional information, as can be seen in Figure 3B, because the confidence intervals of the CAPS scores for REVEL alone and in combination with other filters all overlap. The size of the confidence intervals indicates that the vast majority of variants passing this REVEL threshold also, unsurprisingly, pass the other tools’ thresholds for deleteriousness. Thus, provided that REVEL predictions are trusted by the user, it is possible to discard the rest of the annotations in the report and use the REVEL annotation as evidence of pathogenicity for decision-making.

**Figure 3.**
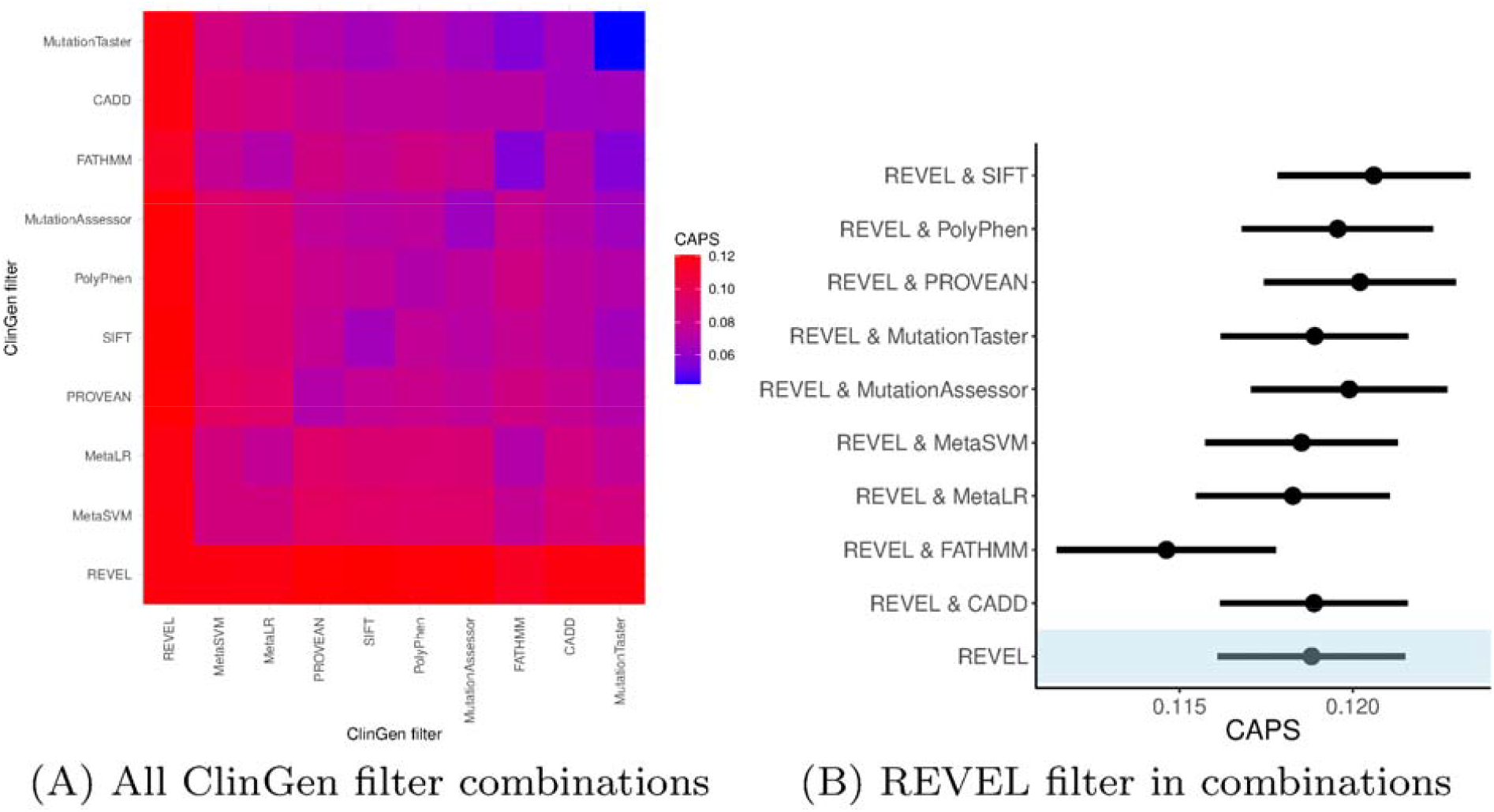
Using CAPS to find key pathogenicity signal in a variant report. ClinGen report was used as an example. CAPS scores of all 1- and 2-filter combinations of threshold-based criteria are shown (panel A), with top-most scores (red) indicating the key annotation that provides evidence of high pathogenicity. The key filter (REVEL, in this case) is also shown alone and in combinations with other filters (panel B). Error bars are 95% binomial confidence intervals. Missense variants only. Variant numbers for each group are shown in Table S5.

We also attempted to recreate the analysis in Figure 2 using the ClinGen results presented above. Specifically, we investigated the possibility of reaching the same high level of deleteriousness achieved by applying the REVEL filter with the other filters from the ClinGen report (Table S2). Although we found that it was not possible to replace the REVEL annotation with any other combination of filters to provide supporting evidence of high pathogenicity, we confirmed that the process of “chaining” filters produced an increase in CAPS with each new filter.

### Meta-benchmarking

Given that ClinVar, a curated database of genetic variation, has traditionally been used for testing and benchmarking of many predictors, we compared the CAPS scores for different categories of ClinVar variants. Figure 4 shows the CAPS scores for ClinVar variants, split using ClinVar’s own star system based on each variant’s review status. Variants reviewed by an expert panel get three stars, variants with two and more submitters with no conflicting interpretations and clear assertion criteria get two stars, variants with one submitter get one star, provided that assertion criteria are available, and other groups get zero stars. For simplicity, in the figure, “Benign” and “Pathogenic” classes include “Likely benign” and “Likely pathogenic” subclasses (the estimates for which are shown in Figures S2 and S3, with additional statistics in Table S3). As shown in Figure 4, benign variants in the expert panel group are no different from the VUSs. In contrast, the groups of variants with fewer stars seem to provide good separation between benign, pathogenic and VUS categories. Besides, Figure S4 shows that the expert panel group is well-calibrated only for LoF variants, but not for missense variants. Based on Figures 4 and S4, we speculate that the ClinVar expert panel may be overly conservative in its assessment of variant pathogenicity. Figure 4 also presents an example of how our approach can be used to assess the average deleteriousness of a curated list of variants. Here we compared our in-house selection of CHD VUSs to all VUSs in ClinVar. We found that the CAPS value for our VUSs was similar to that for ClinVar’s VUSs, with some of our variants being potentially classifiable as either “Benign” or “Pathogenic” due to the large confidence intervals resulting from the small size of our CHD variant set.

**Figure 4.**
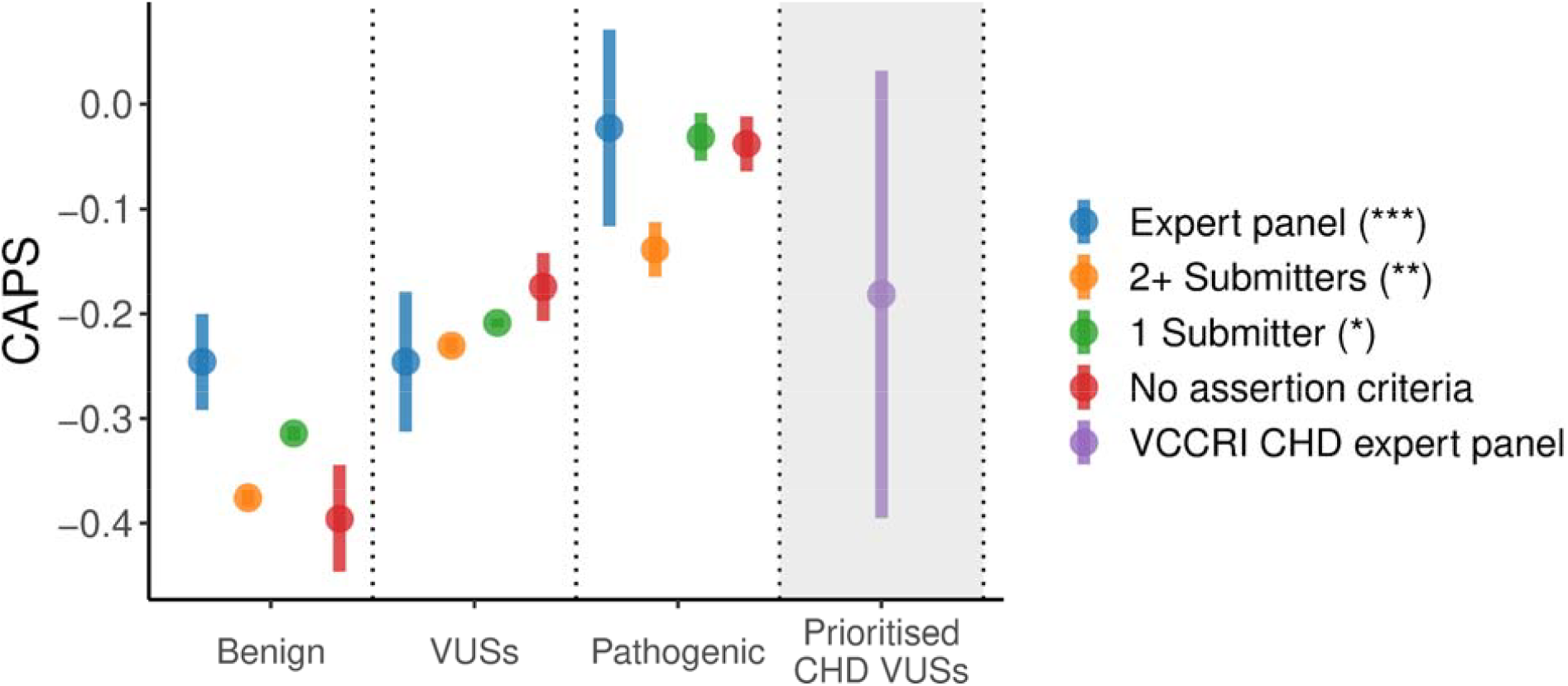
CAPS scores for ClinVar variants split by review status and for an in-house selection of CHD VUSs. “Benign” and “Pathogenic” classes include “Likely benign” and “Likely pathogenic” subclasses respectively. Groups where all variants have a matching ClinVar record have white background; groups where only some variants have a matching ClinVar record have grey background. Error bars are 95% binomial confidence intervals. Variant numbers for each group are shown in Table S6.

CAPS can also be used to show what words like “Damaging” and “Deleterious”, which are often used as classification labels, mean quantitatively. Figure 5 shows the CAPS score for groups of variants that passed the same thresholds from the ClinGen report referred to earlier, together with the labels that each threshold was assigned based on each tool’s classification. The figure highlights that there is substantial variability between the tools in terms of what “Damaging” means. Furthermore, as can be seen in the figure, the definition of “Disease causing” in MutationTaster implies that nearly all missense variants are disease causing, since the average deleteriousness value for the group of variants passing that threshold is close to the genome-wide average. This indicates that most classification labels are poorly calibrated.

**Figure 5.**
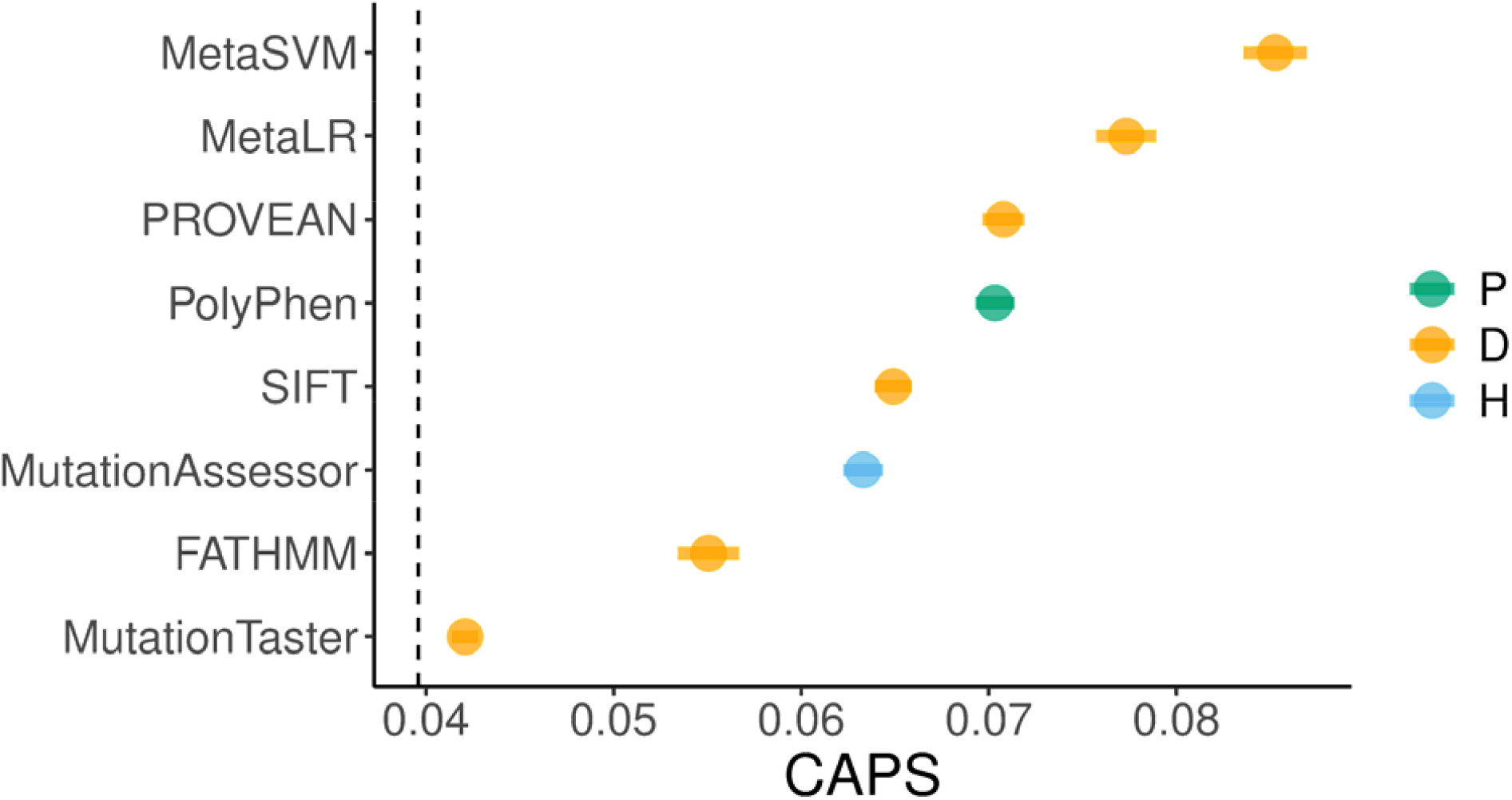
Discrepancy between pathogenicity classification labels and CAPS-informed deleteriousness scores. “D”: “Damaging” (PROVEAN, FATHMM, MetaSVM, MetaLR, SIFT) or “Disease causing” (MutationTaster); “H”: “High” impact (MutationAssessor); “P”: “Possibly damaging” (PolyPhen). Dashed line indicates missense level averaged over all WES variants. Error bars are 95% binomial confidence intervals. Missense variants only. Variant numbers for each group are shown in Table S5.

## Discussion

In this work, we investigated some of the potential applications of population genomics metrics for the benchmarking of commonly used variant pathogenicity predictors and meta-analyses. To this end, we utilised our newly developed CAPS metric, a robust and accurate method that provides estimates of negative selection. Our approach is novel and orthogonal compared to how pathogenicity predictors and their corresponding thresholds have been benchmarked previously.

Using CAPS as our selection-based deleteriousness metric of choice, we found CADD and REVEL to be the best-performing predictors based on their variant separation ability, which allows the user to select variants with high deleteriousness in the absence of score-based thresholds. Moreover, according to our results, REVEL scores appear to be particularly well-calibrated, as REVEL differentiates well between variants even beyond the top 3% point. Therefore, we conclude that finer calibration of score distributions might directly benefit variant filtering and prioritisation pipelines through better variant separation.

These results extend previous studies on the performance of CADD, REVEL and other popular predictors^7,11–14,24^. Importantly, all of these methods have been created and tested based on curated sets of variants, such as those from the ClinVar database. In population-based methods, however, there is no such ascertainment bias (though it may be argued that a similar kind of bias could be the result of, for example, the ethnical composition of population cohorts). Besides, many benchmarking studies suffer the problem of data circularity (i.e., the re-use of data for both training and testing)^2,14,15^. Furthermore, compared to the vast majority of published benchmarking studies, which tend to focus on missense variants^6–8,10,14,24^, our proposed method of utilising a metric of negative selection is more general and agnostic to variant type. This population genomics approach could be used for simple preliminary analyses before employing the more comprehensive methods such as deep mutational scanning (DMS)^30^.

We also demonstrated an effective approach for the assessment and visualisation of the process of “chaining” of filters, which is a common practice in variant prioritisation. Specifically, we showed how CAPS scores change when new filters are added as part of the QC, using some commonly used tool-threshold pairs as an example^27^. We would like to note, however, that in general such combinations of filters may not necessarily be useful, as was argued previously^24^. This is particularly the case when lenient, commonly used thresholds are applied, which, even in combinations, tend to fail at providing sufficient evidence of deleteriousness.

In addition, we present a type of analysis showing how CAPS can be used to find key evidence of deleteriousness in a typical variant report. We found that the threshold for REVEL used in ClinGen reports is the most critical piece of information about the deleteriousness of each variant in these reports.

We note that while our analyses highlight differences in average deleteriousness across variant sets filtered by different annotation thresholds, we do not aim to define optimal thresholds for variant prioritisation. Threshold selection involves a trade-off between sensitivity and specificity, and the appropriate balance depends on the specific application (e.g., clinical versus research settings). Similarly, although REVEL often performed well in our analyses, our goal was not to identify the “best” pathogenicity predictor, but rather to demonstrate how CAPS can be used to evaluate the relative enrichment for deleterious variants across sets filtered by commonly used annotations. The annotations chosen reflect those used in ClinGen variant reports, providing a realistic use case for CAPS in a practical setting.

In line with previously published results exploring the benchmarking of the “ground truth” datasets used for the training and testing of pathogenicity predictors^31^, here we demonstrated how population genomics methods can be used for such meta-benchmarking analyses, using ClinVar as an example. Based on our CAPS-driven assessment of the quality of calibration in ClinVar’s pathogenicity categories, we concluded that there are some discrepancies between the deleteriousness of variants in those categories and our own estimates. We augmented this analysis with a sub-study of CHD VUSs, which we found to be at a deleteriousness level comparable to the average for all ClinVar VUSs.

Overall, this work continues and extends the line of studies exploring population genomics in the context of variant pathogenicity prediction^14,32,33^. However, we emphasise that the use of metrics based on negative selection limits the range of discoverable pathogenic variation. For example, due to the complex evolutionary impact of variants associated with more common complex diseases (such as Alzheimer’s disease or Type 2 diabetes), the resulting negative selection of those variants may not necessarily be strong, and therefore, such variation is unlikely to be fully captured with selection-based metrics. We would also like to note that, despite the above-mentioned criticism of circularity in benchmarking studies, our approach is not completely without circularity either. This is because some of the variant pathogenicity predictors analysed here are based on training sets in which variants are separated by their allele frequencies derived from population data. For example, the PolyPhen predictor is trained using the HumVar variant set, where SNVs with allele frequencies above 1% and no involvement in disease are labelled as “non-damaging”. Similarly, the training of CADD involves population data from BRAVO/TOPMed, to some extent. Despite the fact that our method is based on gnomAD, the allele frequencies may still be correlated.

## Conclusion

Genomic variant annotations are useful if they can aid decision-making or provide insight into biological processes. We believe that annotations which do not assist in these ways only add complexity to genomic analyses, and that with ever-increasing numbers of available annotations, discerning between them has become increasingly important. Population genomics methods, such as CAPS, hold promise for providing an orthogonal, interpretable and efficient way of benchmarking variant pathogenicity predictors and conducting meta-analyses. We would like to emphasise, however, that what we present here is a proof-of-concept analysis and not a comprehensive comparison of pathogenicity predictors, the number of which is constantly growing. We believe that the findings presented in this work will aid analysts in the field of rare disease genomics in their attempts to find new disease-variant associations through efficient variant filtering and prioritisation.

## Supporting information

Supplementary Matereial

## Ethics Declaration

For the CHD data, ethical approval was obtained from the Sydney Children’s Hospital Network Human Research Ethics Committee (approval number HREC/16/SCHN/73). Written informed consent was obtained from all participants. Consent was obtained from a parent or guardian on behalf of any participants under the age of 16.

## References

1. Katsonis P, Wilhelm K, Williams A, Lichtarge O. Genome interpretation using in silico predictors of variant impact. Human Genetics. 2022;141:1549–1577.

2. Karchin R, Radivojac P, O’Donnell-Luria A, Greenblatt MS, Tolstorukov MY, Sonkin D. Improving transparency of computational tools for variant effect prediction. Nature Genetics. 2024;56(7):1324–1326.

3. Jaravine V, Balmford J, Metzger P, Boerries M, Binder H, Boeker M. Annotation of human exome gene variants with consensus pathogenicity. Genes. 2020;11(9).

4. Barbosa P, Ribeiro M, Carmo-Fonseca M, Fonseca A. Clinical significance of genetic variation in hypertrophic cardiomyopathy: Comparison of computational tools to prioritize missense variants. Frontiers in Cardiovascular Medicine. 2022;9.

5. Zhang X, Walsh R, Whiffin N, et al. Disease-specific variant pathogenicity prediction significantly improves variant interpretation in inherited cardiac conditions. Genetics in Medicine. 2021;23:69–79.

6. Tejura M, Fayer S, McEwen AE, Flynn J, Starita LM, Fowler DM. Calibration of variant effect predictors on genome-wide data masks heterogeneous performance across genes. The American Journal of Human Genetics. 2024;111:2031–2043.

7. Tian Y, Pesaran T, Chamberlin A, et al. REVEL and BayesDel outperform other in silico meta-predictors for clinical variant classification. Scientific Reports. 2019;9.

8. Zaucha J, Heinzinger M, Tarnovskaya S, Rost B, Frishman D. Family-specific analysis of variant pathogenicity prediction tools. NAR Genomics and Bioinformatics. 2020;2(2):qaa014.

9. Accetturo M, Bartolomeo N, Stella A. In-silico analysis of NF1 missense variants in ClinVar: Translating variant predictions into variant interpretation and classification. International Journal of Molecular Sciences. 2020;21(3).

10. Qorri E, Takács B, Gráf A, et al. A comprehensive evaluation of the performance of prediction algorithms on clinically relevant missense variants. International Journal of Molecular Sciences. 2022;23(14).

11. Cubuk C, Garrett A, Choi S, et al. Clinical likelihood ratios and balanced accuracy for 44 in silico tools against multiple large-scale functional assays of cancer susceptibility genes. Genetics in Medicine. 2021;23(11):2096–2104.

12. Gunning AC, Fryer V, Fasham J, et al. Assessing performance of pathogenicity predictors using clinically relevant variant datasets. Journal of Medical Genetics. 2021;58(8):547–555.

13. Wang D, Li J, Wang Y, Wang E. A comparison on predicting functional impact of genomic variants. NAR Genomics and Bioinformatics. 2022;4(1).

14. Tabet DR, Kuang D, Lancaster MC, et al. Benchmarking computational variant effect predictors by their ability to infer human traits. Genome Biology. 2024;25(1):172.

15. Livesey BJ, Marsh JA. Interpreting protein variant effects with computational predictors and deep mutational scanning. Disease Models & Mechanisms. 2022;15(6):dmm049510.

16. Shah N, Hou YCC, Yu HC, et al. Identification of misclassified ClinVar variants via disease population prevalence. 2018;102:609–619.

17. Karczewski KJ, Francioli LC, Tiao G, et al. The mutational constraint spectrum quantified from variation in 141,456 humans. Nature. 2020;581(7809):434–443.

18. Gudmundsson S, Singer-Berk M, Watts NA, et al. Variant interpretation using population databases: Lessons from gnomAD. Human Mutation. 2022;43(8):1012–1030.

19. Gudkov M, Thibaut L, Giannoulatou E. Context-adjusted proportion of singletons (CAPS): A novel metric for assessing negative selection in the human genome. NAR Genomics and Bioinformatics. 2024;6(3).

20. Lek M, Karczewski KJ, Minikel EV, et al. Analysis of protein-coding genetic variation in 60,706 humans. Nature. 2016;536(7616):285–291.

21. Harpak A, Bhaskar A, Pritchard JK. Mutation rate variation is a primary determinant of the distribution of allele frequencies in humans. PLOS Genetics. 2016;12(12):1–22.

22. Samocha KE, Robinson EB, Sanders SJ, et al. A framework for the interpretation of de novo mutation in human disease. Nature Genetics. 2014;46(9):944–950.

23. Short PJ, McRae JF, Gallone G, et al. De novo mutations in regulatory elements in neurodevelopmental disorders. Nature. 2018;555:611–616.

24. Pejaver V, Byrne AB, Feng BJ, et al. Calibration of computational tools for missense variant pathogenicity classification and ClinGen recommendations for PP3/BP4 criteria. The American Journal of Human Genetics. 2022;109:2163–2177.

25. Liu X, Li C, Mou C, Dong Y, Tu Y. dbNSFP v4: A comprehensive database of transcript-specific functional predictions and annotations for human nonsynonymous and splice-site SNVs. Genome Medicine. 2020;12.

26. Alankarage D, Ip E, Szot JO, et al. Identification of clinically actionable variants from genome sequencing of families with congenital heart disease. Genetics in Medicine. 2019;21(5):1111–1120.

27. Jagadeesh KA, Wenger AM, Berger MJ, et al. M-CAP eliminates a majority of variants of uncertain significance in clinical exomes at high sensitivity. Nature Genetics. 2016;48:1581–1586.

28. Aoki E, Manabe N, Ohno S, et al. Predicting the pathogenicity of missense variants based on protein instability to support diagnosis of patients with novel variants of ARSL. Molecular Genetics and Metabolism Reports. 2023;37:101016.

29. Canning AJ, Viggiano S, Fernandez-Zapico ME, Cosgrove MS. Parallel functional annotation of cancer-associated missense mutations in histone methyltransferases. Scientific reports. 2022;12(1):18487.

30. Livesey BJ, Marsh JA. Updated benchmarking of variant effect predictors using deep mutational scanning. Molecular Systems Biology. 2023;19(8):e11474.

31. Sarkar A, Yang Y, Vihinen M. Variation benchmark datasets: update, criteria, quality and applications. Database. 2020;2020:baz117.

32. Niroula A, Vihinen M. How good are pathogenicity predictors in detecting benign variants? PLOS Computational Biology. 2019;15(2):1–17.

33. Cabrera-Alarcon JL, Martinez JG, Enríquez JA, Sánchez-Cabo F. Variant pathogenic prediction by locus variability: The importance of the current picture of evolution. European Journal of Human Genetics. 2022;30:555–559.

