## Supplementary Matereial for "Benchmarking of variant pathogenicity prediction methods using a population genetics approach"

### Supplemental Note 1

"Chaining" of filters refers to the process of incremental addition of quality control (QC) filters applied to a variant set. For example, filter $F_1$ can be "REVEL>0.75". Naturally, any two chains made of the same QC filters will produce identical variant sets, regardless of the order in which these filters are applied in each of the two chains.

To test all possible combinations of filters $F_1, F_2, \dots, F_N$ for changes in the CAPS value, we used the following algorithm:

1. Make a list of all $N$ filters for the analysis
2. Select maximum chain length $M$ (i.e. the maximum number of filters to be included in a single chain, $M < N$)
3. For chain length from 1 to $M$:
    1. Generate all possible combinations of filters for the current length
    2. Apply each of the combinations, thus generating a new variant set
    3. Calculate CAPS on each of the new variant sets
    4. Select the maximum CAPS value for the current iteration

Hence, chains of 1 will be $(F_1), (F_2), (F_3), \dots$; chains of 2 will be $(F_1 \times F_2), (F_1 \times F_3), (F_2 \times F_3), \dots$, and so on.

### Supplemental Figures

ClinGen Predictors

| Source | Score Range | Score | Impact Threshold | Prediction |
|---|---|---|---|---|
| REVEL (meta-predictor) | 0 to 1 | 0.782 | >0.75 | higher score = higher pathogenicity |

Other Predictors

| Source | Score Range | Score | Impact Threshold | Prediction |
|---|---|---|---|---|
| SIFT | -- | 0.005, 0.005 | <0.049 | D, D |
| PolyPhen2-HDIV | 0 to 1 | 0.995, 0.995 | -- | D, D |
| PolyPhen2-HVAR | 0 to 1 | 0.893, 0.893 | >0.447 | P, P |
| LRT | 0 to 1 | 0.004033 | -- | N |
| MutationTaster | 0 to 1 | 0.997549, 0.997549 | >0.5 | D, D |
| MutationAssessor | -0.5135 to 6.49 | 3.965, 3.965 | >1.935 | H, H |
| FATHMM | -16.13 to 10.64 | -3.31, -3.31 | <-1.51 | D, D |
| PROVEAN | -14 to +14 | -4.02, -4.02 | <-2.49 | D, D |
| MetaSVM (meta-predictor) | -2 to +3 | 0.9853 | >0 | D |
| MetaLR (meta-predictor) | 0 to 1 | 0.8946 | >0.5 | D |

**Figure S1**. Example of a ClinGen report for a missense variant as shown in ClinGen's Variant Curation Interface. The thresholds displayed are relevant as of 2024/09/25.

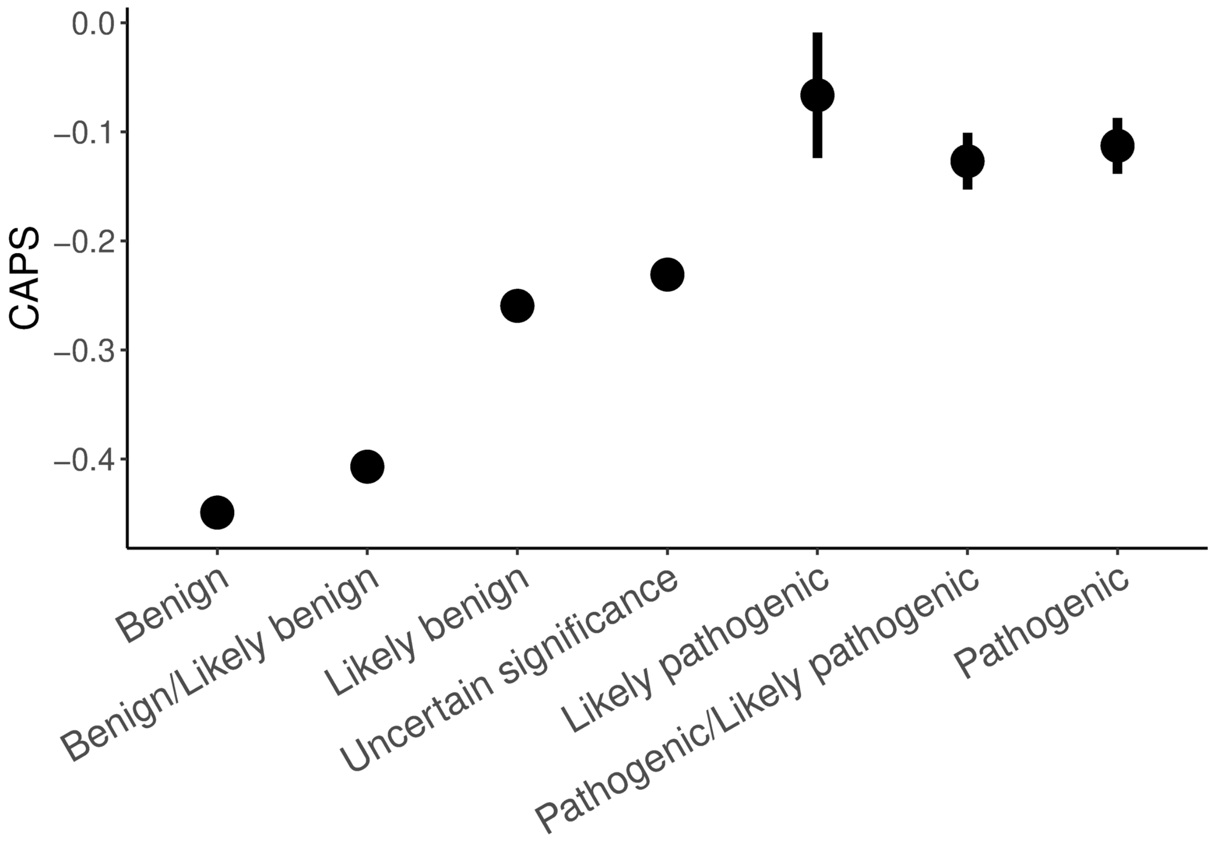


**Figure S2**. CAPS scores for groups of variants stratified by ClinVar pathogenicity. Only germline variants with at least two independent submitters and no conflicting interpretations were included.

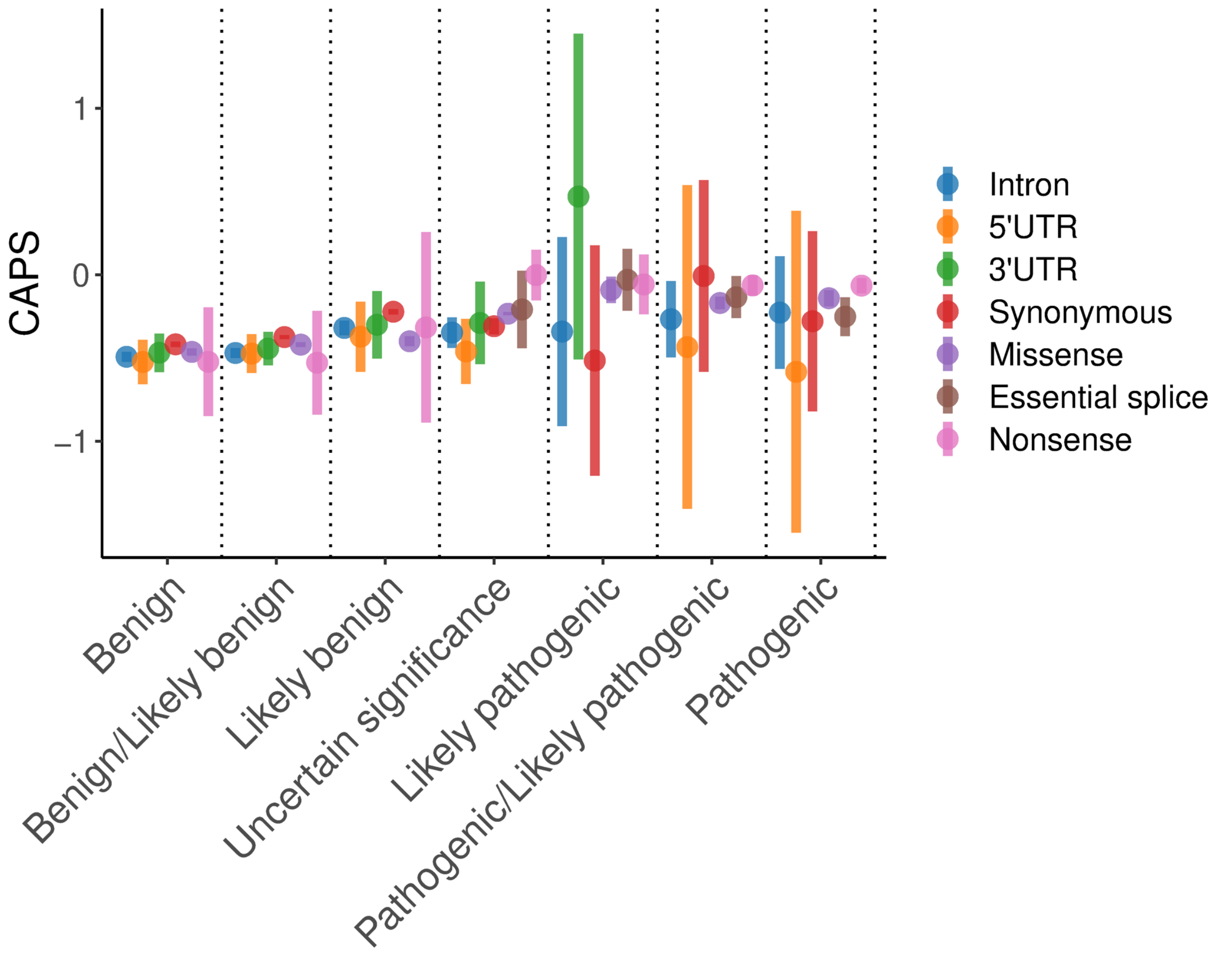


**Figure S3**. CAPS scores for groups of variants stratified by ClinVar pathogenicity and variant type. Only germline variants with at least two independent submitters and no conflicting interpretations were included.

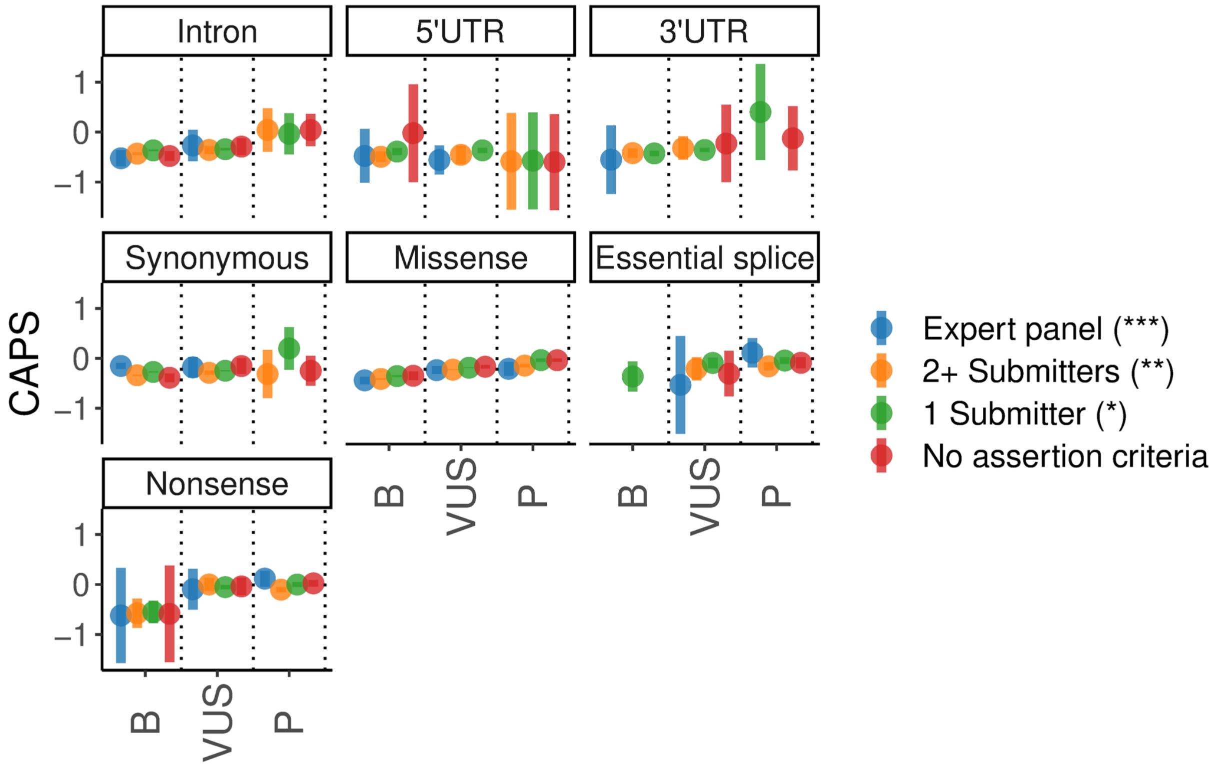


**Figure S4**. CAPS scores for all ClinVar variants split by review status and variant type. "Benign" ("B") and "Pathogenic" ("P") categories include "Likely benign" and "Likely pathogenic" categories respectively.

### Supplemental Tables

**Table S1**. The total number of variation observed in each variant class after preprocessing.

| **Class** | **Total** |
|---|---|
| Intron | 2436006 |
| 5’UTR | 108405 |
| 3’UTR | 160419 |
| Synonymous | 2173989 |
| Missense | 4547687 |
| Essential splice | 65907 |
| Nonsense | 132974 |

**Table S2**. Change in CAPS scores during the process of finding supporting evidence for the key pathogenicity signal in a variant report, using all reference-free combinations. ClinGen report was used as an example. REVEL score of 0.118 was used as a reference. Chain length of 1 indicates one filter (based on one tool-threshold pair), chain length of 2 indicates two filters in combination, etc. Missense variants only.

| **Chain length** | **Max CAPS** | **CAPS distance to reference** | **CAPS increase (percent)** |
|---|---|---|---|
| 1 | 0.0849 | 0.0336 | NA |
| 2 | 0.0959 | 0.0226 | 32.6 |
| 3 | 0.0994 | 0.0190 | 15.7 |
| 4 | 0.102 | 0.0169 | 11.3 |
| 5 | 0.103 | 0.0158 | 6.19 |
| 6 | 0.103 | 0.0151 | 4.69 |
| 7 | 0.104 | 0.0146 | 3.43 |
| 8 | 0.104 | 0.0145 | 0.511 |
| 9 (all) | 0.101 | 0.0172 | -18.7 |

**Table S3**. Per-group statistics for ClinVar-annotated germline variants. Subsets containing all variants (ALL) and variants with at least two independent submitters and no conflicting interpretations (high-confidence, HC) are compared.

| **Clinical significance** | **Total (ALL)** | **Proportion singletons (ALL)** | **Total (HC set)** | **Proportion singletons (HC set)** |
|---|---|---|---|---|
| Benign | 17278 | 0.006 | 6409 | 0.000 |
| Benign/Likely benign | 11543 | 0.002 | 11508 | 0.002 |
| Likely benign | 40264 | 0.149 | 7083 | 0.141 |
| Uncertain significance | 83162 | 0.199 | 17471 | 0.171 |
| Pathogenic/Likely pathogenic | 1505 | 0.214 | 1442 | 0.212 |
| Likely pathogenic | 3252 | 0.366 | 365 | 0.315 |
| Pathogenic | 7708 | 0.329 | 1413 | 0.220 |

**Table S4**. Variant numbers for Figure 2 illustrating the effect of using stringent thresholds and "chaining" of filters on resulting CAPS scores.

| **Group** | **Thresholds** | **Number of variants** |
|---|---|---|
| SIFT | Less stringent | 3265568 |
| CADD | Less stringent | 3244444 |
| PolyPhen | Less stringent | 2722536 |
| All 3 | Less stringent | 2483104 |
| SIFT | More stringent | 1858218 |
| CADD | More stringent | 2349036 |
| PolyPhen | More stringent | 1264886 |
| All 3 | More stringent | 1005175 |

**Table S5**. Variant numbers for Figure 3 and Figure 5, illustrating CAPS scores for individual filters and combinations of filters.

| Group | Number of variants |
|---|---|
| REVEL | 240971 |
| SIFT | 1848485 |
| PolyPhen | 1691426 |
| MutationTaster | 3706267 |
| MutationAssessor | 1602851 |
| FATHMM | 669160 |
| PROVEAN | 1458721 |
| MetaSVM | 628402 |
| MetaLR | 690356 |
| CADD | 2458634 |
| REVEL & SIFT | 228382 |
| REVEL & PolyPhen | 231264 |
| REVEL & MutationTaster | 240908 |
| REVEL & MutationAssessor | 220665 |
| REVEL & FATHMM | 175599 |
| REVEL & PROVEAN | 230387 |
| REVEL & MetaSVM | 229331 |
| REVEL & MetaLR | 225766 |

| Group | Number of variants |
|---|---|
| REVEL & CADD | 240816 |
| SIFT & PolyPhen | 1308310 |
| SIFT & MutationTaster | 1846668 |
| SIFT & MutationAssessor | 1217062 |
| SIFT & FATHMM | 377341 |
| SIFT & PROVEAN | 1188928 |
| SIFT & MetaSVM | 507308 |
| SIFT & MetaLR | 507380 |
| SIFT & CADD | 1596622 |
| PolyPhen & MutationTaster | 1689684 |
| PolyPhen & MutationAssessor | 1173121 |
| PolyPhen & FATHMM | 355732 |
| PolyPhen & PROVEAN | 1115918 |
| PolyPhen & MetaSVM | 518183 |
| PolyPhen & MetaLR | 516255 |
| PolyPhen & CADD | 1583973 |
| MutationTaster & MutationAssessor | 1601038 |
| MutationTaster & FATHMM | 668408 |
| MutationTaster & PROVEAN | 1457185 |
| MutationTaster & MetaSVM | 628109 |

| Group | Number of variants |
|---|---|
| MutationTaster & MetaLR | 690037 |
| MutationTaster & CADD | 2455747 |
| MutationAssessor & FATHMM | 331150 |
| MutationAssessor & PROVEAN | 1069397 |
| MutationAssessor & MetaSVM | 477210 |
| MutationAssessor & MetaLR | 476010 |
| MutationAssessor & CADD | 1389928 |
| FATHMM & PROVEAN | 324846 |
| FATHMM & MetaSVM | 422472 |
| FATHMM & MetaLR | 500074 |
| FATHMM & CADD | 502084 |
| PROVEAN & MetaSVM | 464763 |
| PROVEAN & MetaLR | 458839 |
| PROVEAN & CADD | 1308087 |
| MetaSVM & MetaLR | 587016 |
| MetaSVM & CADD | 600186 |
| MetaLR & CADD | 617033 |

**Table S6**. Variant numbers for Figure 4 illustrating CAPS scores for ClinVar variants split by review status and for an in-house selection of CHD VUSs.

| **Group** | **Review status** | **Number of variants** |
|---|---|---|
| CHD VUS | reviewed by VCCRI CHD expert panel | 31 |
| Pathogenic | reviewed by expert panel | 205 |
| VUS | reviewed by expert panel | 340 |
| Benign | reviewed by expert panel | 757 |
| Pathogenic | criteria provided, multiple submitters, no conflicts | 2063 |
| Benign | criteria provided, multiple submitters, no conflicts | 14941 |
| VUS | criteria provided, multiple submitters, no conflicts | 22175 |
| Pathogenic | criteria provided, single submitter | 3062 |
| Benign | criteria provided, single submitter | 24652 |
| VUS | criteria provided, single submitter | 77699 |
| Benign | no assertion criteria provided | 392 |
| VUS | no assertion criteria provided | 1342 |
| Pathogenic | no assertion criteria provided | 2309 |
